# A Dual Color Pax7 and Myf5 In Vivo Reporter to Investigate Muscle Stem Cell Heterogeneity in Regeneration and Aging

**DOI:** 10.1101/2023.06.19.545587

**Authors:** Sara Ancel, Joris Michaud, Federico Sizzano, Loic Tauzin, Manuel Oliveira, Eugenia Migliavacca, Gabriele Dammone, Sonia Karaz, José L Sánchez-García, Sylviane Metairon, Guillaume Jacot, C. Florian Bentzinger, Jérôme N. Feige, Pascal Stuelsatz

## Abstract

Increasing evidence suggests heterogeneity in the muscle stem cell (MuSC) pool. In particular, a rare subset of Pax7 positive MuSCs that has never expressed the myogenic regulatory factor Myf5 has enhanced self-renewal and engraftment characteristics. However, the scarcity and limited availability of protein markers make the characterization of these cells challenging. We describe the generation of StemRep reporter mice allowing to monitor Pax7 and Myf5 protein based on equimolar levels of dual nuclear fluorescence. High levels of Pax7 protein and low levels of Myf5 delineate a deeply quiescent MuSC subpopulation with distinct molecular signatures and dynamics of activation, proliferation, and commitment. Aging decreases the number of these cells, and skews the MuSC pool towards Myf5-^High^ cells with impaired quiescence. Altogether, we describe a novel deeply quiescent MuSC subpopulation whose maintenance is impaired in old muscles, and establish the StemRep line as a versatile tool to study quiescence and MuSC heterogeneity.

## Introduction

Muscle stem cells (MuSCs) are tissue-resident stem cells that are essential for growth, maintenance, and repair of adult skeletal muscle throughout life (Bachman et al., 2018; Englund et al., 2020; Lepper et al., 2011; Sambasivan et al., 2011). Under homeostatic conditions, adult MuSCs reside in a quiescent state in a niche compartment located between the basal lamina and the muscle fiber plasma membrane (Dumont et al., 2015; Mauro, 1961; Schüler et al., 2022). The long-term regenerative potential of skeletal muscle relies on the ability of MuSCs to maintain, exit, and re-enter quiescence, a reversible cell cycle-arrested state allowing the cells to respond to different environmental signals (Ancel et al., 2021; van Velthoven and Rando, 2019). Age-associated regenerative failure of skeletal muscle is governed by defective cell-intrinsic mechanisms controlling features such as quiescence and activation, as well as by aberrant niche signals in the MuSC microenvironment (Ancel et al., 2021; Mashinchian et al., 2018< Sousa-Victor et al., 2022). Despite considerable recent success in characterizing these mechanisms, important unresolved questions about the balance of intrinsic and extrinsic aging and how it affects the stem cell pool remain (Ancel et al., 2021; Sousa-Victor et al., 2022).

MuSCs express the paired-box transcription factor Pax7, a canonical myogenic marker that represses genes involved in differentiation, which is essential for proper muscle regeneration (Ancel et al., 2021; van Velthoven and Rando, 2019; von Maltzahn et al., 2013). Once they break quiescence, MuSCs upregulate myogenic regulatory factors (MRFs), a family of transcription factors associated with lineage progression and commitment to differentiation (Olguin and Olwin, 2004; Sambasivan et al., 2011; Seale et al., 2000). Activated MuSCs contain high levels of Myf5 and MyoD, and subsequently progress towards terminal differentiation by down-regulating Pax7, Myf5 and MyoD, while increasing the expression of Myogenin (Myog) and MRF4 (Schmidt et al., 2019; Zammit, 2017).

Compelling evidence points toward molecular heterogeneity in the MuSC pool. Certain MuSC subpopulations show differential regulation of cell cycle progression, lineage commitment, self-renewal, repopulation of the niche, and the ability to resist environmental stress (Ancel et al., 2021). High Pax7 levels are associated with deeply quiescent MuSCs that are slower to enter the cell cycle, have a lower metabolic activity, and greater ability to engraft in serial transplantations (Rocheteau et al., 2012). Similarly, expression of the surface markers CD34, CD9, and CD104 has been linked to differential capabilities of MuSCs to repopulate the niche after transplantation (Beauchamp et al., 2000; García-Prat et al., 2020; Porpiglia et al., 2017). Pax3 expression has been associated with a subset of MuSCs in deeper quiescence that show a higher resistance and survival advantages in response to environmental stresses (Der Vartanian et al., 2019; Scaramozza et al., 2019). Apart from the downregulation of stemness genes involved in maintaining quiescence, myogenic commitment to differentiation of MuSC subpopulations has been shown to depend on MRFs. In particular, Myf5 appears to play a central role in the epigenic specification and committment of MuSCs towards differentiation (Beauchamp et al., 2000; Kuang et al., 2007). Using Myf5-Cre;ROSA26-YFP mice, it has been demonstrated that MuSCs that are lineage negative for Myf5 self-renew through asymmetric divisions generating one committed daughter cell while maintaining a mother “satellite stem cell” (Kuang et al., 2007). Myf5 lineage negative cells also show a higher engraftment and repopulation potential than lineage positive cells. Supporting these findings, it has been shown that Pax7 positive MuSCs that are haplo-insufficient for Myf5 show an increased ability to self-renew and perform better in transplantation assays than wild-type controls (Gayraud-Morel et al., 2012). Altogether, these observations suggest that the stem cell pool in skeletal muscle is maintained by a MuSC subpopulation with superior stem cell characteristics. Importantly, given the deleterious effects of aging on the regenerative capacity of skeletal muscle and the MuSC pool, it is conceivable that this stem cell subpopulation is disrupted in aged muscle tissue.

Much of our current understanding of the Myf5 negative MuSC population is based on lineage tracing, which does not allow for conclusions regarding the real-time dynamics of this transcription factor in MuSCs. In addition, untranslated Myf5 mRNA is sequestered into messenger ribonucleoprotein (mRNP) granules, which makes the prediction of protein levels using gene expression reporters challenging (Crist et al., 2012). Moreover, knock-in reporter alleles such as the Myf5^LacZ^ or Myf5^GFP^ lines may be confounded by Myf5 haploinsufficiency leading to altered MuSC function (Beauchamp et al., 2000; Kassar-Duchossoy et al., 2004; Tajbakhsh et al., 1996). Lastly, antibodies for the faithful detection of Myf5 protein in quiescent or activated MuSCs have been difficult to validate. For these reasons, the regulation of Myf5 protein in adult Pax7 positive MuSCs and its potential implication in the aging process remain unclear.

To resolve this long-standing problem and study Myf5 protein levels in adult Pax7 positive MuSCs *in vivo* during regeneration and aging, we generated a dual color reporter mouse line that we named StemRep. To this end, we linked the last exon of the *PAX7* gene to an in-frame T2A peptide and a nuclear targeted zsGreen1 fluorescent protein and used the same strategy to connect a nuclear targeted far-red fluorescent E2-Crimson to *MYF5*. In double heterozygous StemRep mice harboring these two alleles, Pax7 and Myf5 are expressed as a single transcript containing the uninterrupted coding frame linked to the nuclear targeted fluorescent reporter proteins by T2A. Thus, post-translational cleavage of T2A in StemRep mice leads to the induction of dual nuclear fluorescence at equimolecular levels with Myf5 and Pax7 protein. Here, we demonstrate that the StemRep line allows for accurate *in vivo* assessment of Myf5 and Pax7 protein levels in MuSCs and enables efficient flow cytometry isolation of different stem cell subpopulations. We discovered that MuSCs containing high levels of Pax7 (Pax7^High^) and low levels of Myf5 (Myf5^Low^) protein display a clear propensity towards quiescence, including retention of Pax7 expression and a delay in MyoD induction after activation, slower cell-cycle entry, and a lower ability for clonal expansion. Gene expression profiling revealed that Myf5^High^ cells are primed for activation, cell cycle entry, and are committed to myogenic differentiation. Aging induces a decline in the number of total Pax7 positive cells and induces skewing towards a higher proportion of Myf5^High^ MuSCs. Altogether, using StemRep mice, we demonstrate for the first time that low levels of Myf5 protein in Pax7 positive cells represents a key feature of deep stem cell quiescence and provide the field with a novel tool for the assessment of heterogeneity in the adult MuSC pool under different physiological conditions.

## Results

### Generation and validation of StemRep mice

To produce dual Pax7 and Myf5 protein reporter (StemRep) mice, two separate transgenic lines were generated and later intercrossed. Briefly, a nuclear localization signal (NLS) fused to Zoanthus sp. Green fluorescent protein (NLS-ZsGreen1) and the DsRed-derived E2-Crimson far-red fluorescent protein(NLS-E2Crimson) were inserted in frame downstream of the last exon of Pax7 and Myf5 genes, preserving the entire coding region of both genes (Matz et al., 1999; Strack et al., 2009) (**Figures 1A,B**). Using this approach, Pax7 and Myf5 are co-translated with their respective fusion proteins to produce equimolar amounts of the reporter proteins. Both reporters were preceded by a self-cleaving T2A peptide sequence to induce post-translational cleavage of the reporter and production of an independent functional Pax7 or Myf5 protein. Successful targeting of both constructs was confirmed by PCR (**Figure S1A**) and double homozygous animals were selected for subsequent applications based on greater signal-to-noise ratio of ZsGreen1 and E2Crimson fluorescence (**Figure S1B**). No apparent effects of the construct were observed on hindlimb muscle weight when StemRep mice were compared to WT animals at different ages (**Figure S1C**).

**Figure 1:**
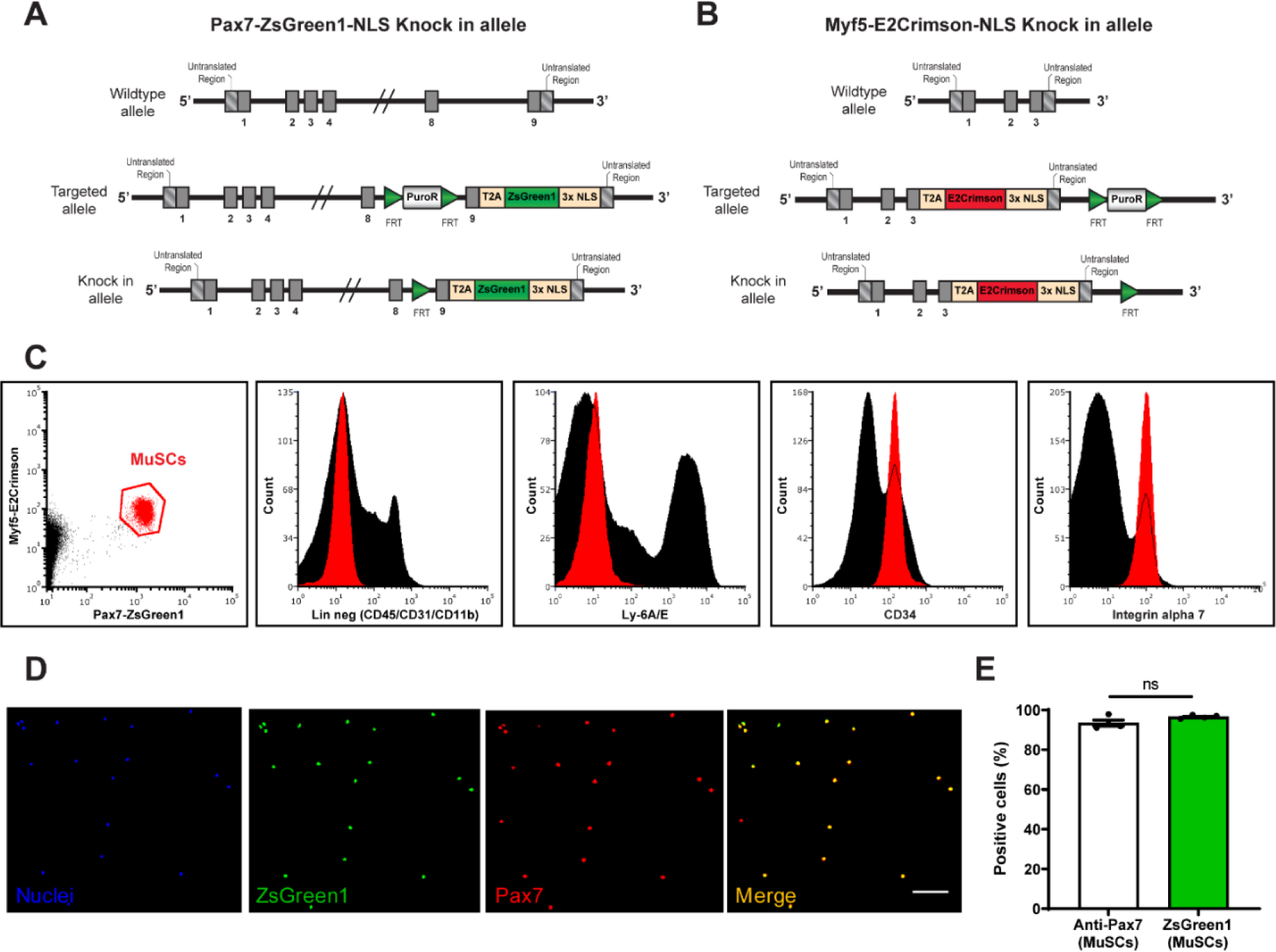
Generation and characterization of StemRep mice. (**A**,**B**) Genetic constructs of (A) Pax7-ZsGreen1-NLS and (B) Myf5-E2Crimson-NLS. Representation of the wild-type allele (top), targeted allele (middle), and knock-in allele after recombination (bottom). (**C**) Representative flow cytometry plots of MuSCs (red) derived from StemRep mice and the associated CD11b-/CD31-/CD45-/Sca1-/CD34+/Itga7+ antigen signature. (**D**,**E**) Post-sort representative images of cytospin preparations (D) and quantification (E) of MuSCs isolated based on nuclear ZsGreen1 fluorescence that stained positive using antibodies for Pax7. ZsGreen1 reporter, green; Pax7, red; Nuclei (Hoechst), blue. Scale bar = 50µm. n = 4 mice. Data presented are means ± s.e.m. ns = not significant in a two-tailed unpaired Student’s t-test (E).

We next analyzed the expression of MuSC-specific surface markers in ZsGreen1-positive cells of StemRep mice (Maesner et al., 2016). Pax7-ZsGreen1 cells were negative for lineage markers (CD31, CD11b, CD45) and Sca1, but positive for CD34 and α7-integrin, consistent with the expected surface antigen signature of MuSCs (**Figure 1C** and **Figure S1D**). Conversely, all CD34 and α7-integrin-positive MuSCs expressed Pax7-ZsGreen1 (**Figure S1E)**. Post-sort analysis using antibody staining showed an enrichment of Pax7 positive cells in the ZsGreen1 fraction (**Figures 1D,E**). To further confirm the specificity of the StemRep Pax7 and Myf5 dual reporter alleles in labeling MuSCs, different muscles and tissues were analyzed by flow cytometry. While cells isolated from hindlimb, extraocular, and diaphragm muscles emitted in the ZsGreen1 and E2Crimson channels, no fluorescence was detected in either brain, liver, or kidney (**Figure S1F**). Endogenous ZsGreen1 fluorescence and immunostaining of cryosections using DsRed antibodies against E2Crimson revealed robust nuclear fluorescence in the *tibialis anterior* (TA) of StemRep mice, while no signal was detected in the brain, kidney, or liver (**Figure S1G**). Altogether, these data demonstrate that Pax7 and Myf5 dependent dual nuclear fluorescence in StemRep mice is specific to MuSCs.

To further characterize MuSCs in StemRep mice, we isolated ZsGreen1-positive cells, cultured them in high-serum conditions, and live-imaged them every 12 hours over a course of 4 days (**Figures S2A-C**) (Bentzinger et al., 2012). ZsGreen1-positive cells were able to differentiate and formed elongated myocytes as their confluence increased (**Figures S2A,B**). This was accompanied by a progressive loss of Pax7 expression as read out by ZsGreen1 fluorescence (**Figures S2A** and **C**). Thus, nuclear ZsGreen1 fluorescence in MuSCs isolated from StemRep mice dynamically reflects Pax7 protein levels.

### Myf5-E2Crimson levels identify a Myf5^High^ and Myf5^Low^ MuSC population

Taking advantage of the StemRep model, we evaluated the capability of the E2Crimson reporter to discriminate Myf5 protein levels. Two subpopulations were isolated by FACS at opposite ends of the E2Crimson spectrum and labeled as Myf5^High^ and Myf5^Low^ cells (**Figure 2A**). Re-analysis of sorted cells by flow cytometry confirmed that the isolated subpopulations consistently fell within the corresponding gates (**Figures S3A,B**). Fluorescence imaging after sorting revealed that the majority of isolated Myf5^Low^ cells were positive for ZsGreen1 fluorescence only (**Figures 2B,C**). In contrast, in only 7% of the sorted Myf5^High^ population no live E2-Crimson fluorescence was detectable while 90% contained both fluorescent markers and 3% exhibited exclusive E2-Crimson fluorescence (**Figures 2B,C**). Average intensity quantification of Myf5-E2Crimson fluorescence in each population further verified that the gating strategy robustly discriminated Myf5 subpopulations (**Figures 2D,E**). We observed that ZsGreen1 fluorescence tracking Pax7 expression was higher in Myf5^Low^ compared to Myf5^High^ cells (**Figure 2F**). In summary, these data demonstrate that the Myf5^Low^ population in StemRep mice contains higher levels of Pax7, which supports the notion that they represent uncommitted MuSCs with superior stem cell characteristics compared to Myf5^High^ cells.

**Figure 2:**
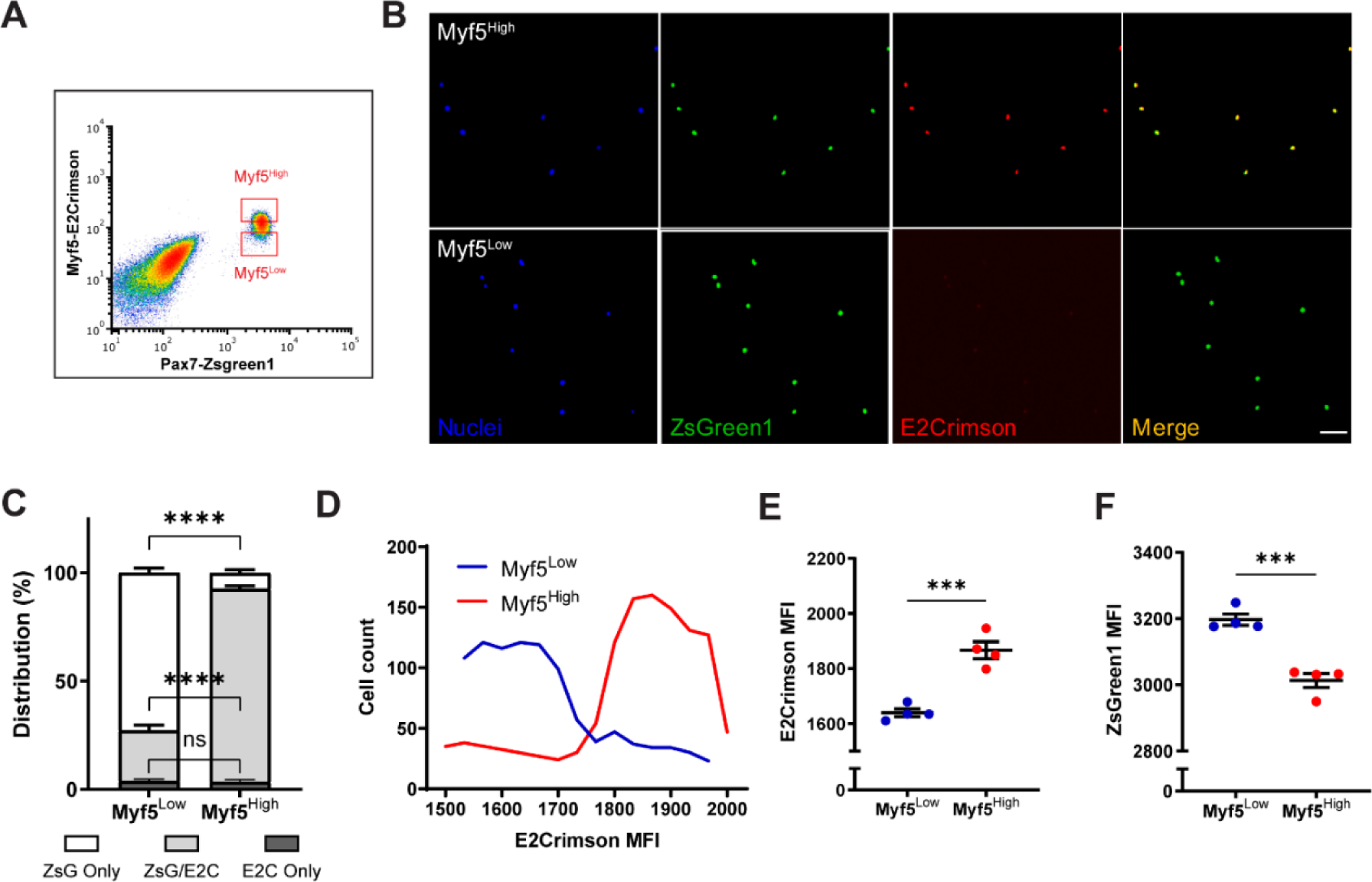
Comparative characterization of freshly isolated Myf5^High^ and Myf5^Low^ MuSCs. (**A**) Flow cytometry-based isolation strategy for Myf5^High^ and Myf5^Low^ MuSCs. (**B**) Representative images of Myf5^High^ and Myf5^Low^ cells in cytospin preparations immediately after sorting. Scale bar, 100µm. (**C**) Histogram showing the proportions of cells positive for either Pax7-ZsGreen1 or Myf5-E2Crimson, or both in isolated Myf5^High^ and Myf5^Low^ fractions. (**D**) Distribution of Myf5^High^ and Myf5^Low^ cells according to E2Crimson intensity following isolation. A total of 1000 cells per subpopulation were analyzed. (**E-F**) Average intensity of E2Crimson (E) and ZsGreen1 (F) fluorescence in each subpopulation. n = 4 mice. Data presented are means ± s.e.m.; ***p<0.001, ****p<0.0001 in a two-tailed unpaired Student’s t-test (C, E, F).

### *In vivo* dynamics of Myf5 subpopulations during muscle regeneration and aging

Because of the central role of MuSCs during skeletal muscle regeneration, we capitalized on the StemRep model to dynamically monitor Myf5 subpopulations at different stages after activation. We first isolated MuSCs after cardiotoxin injury and analyzed different timepoints by flow cytometry. Consistent with a downregulation of both Pax7 and Myf5 during myogenic progression at early timepoints after injury (Bentzinger et al., 2012; Schmidt et al., 2019), the fluorescence intensity of both reporters at 3 and 7 days post injury (dpi) markedly decreased (**Figures 3A,B** and **Figure S4A**). ZsGreen1 and E2Crimson fluorescence was progressively restored to basal levels as MuSCs re-entered quiescence after repair in later stages at 14 and 21 dpi. Immunostaining quantification using ZsGreen1 endogenous fluorescence and DsRed antibody against E2Crimson showed that the number of myogenic progenitors increased over 20-fold at 3 dpi before returning to homeostatic levels around 21 dpi (**Figures 3C,D**). These data demonstrate that StemRep mice allow to monitor of Pax7 and Myf5 protein levels in MuSCs and reveal a Myf5^Low^ population that is present under homeostatic conditions and persists across all stages of skeletal muscle regeneration.

**Figure 3:**
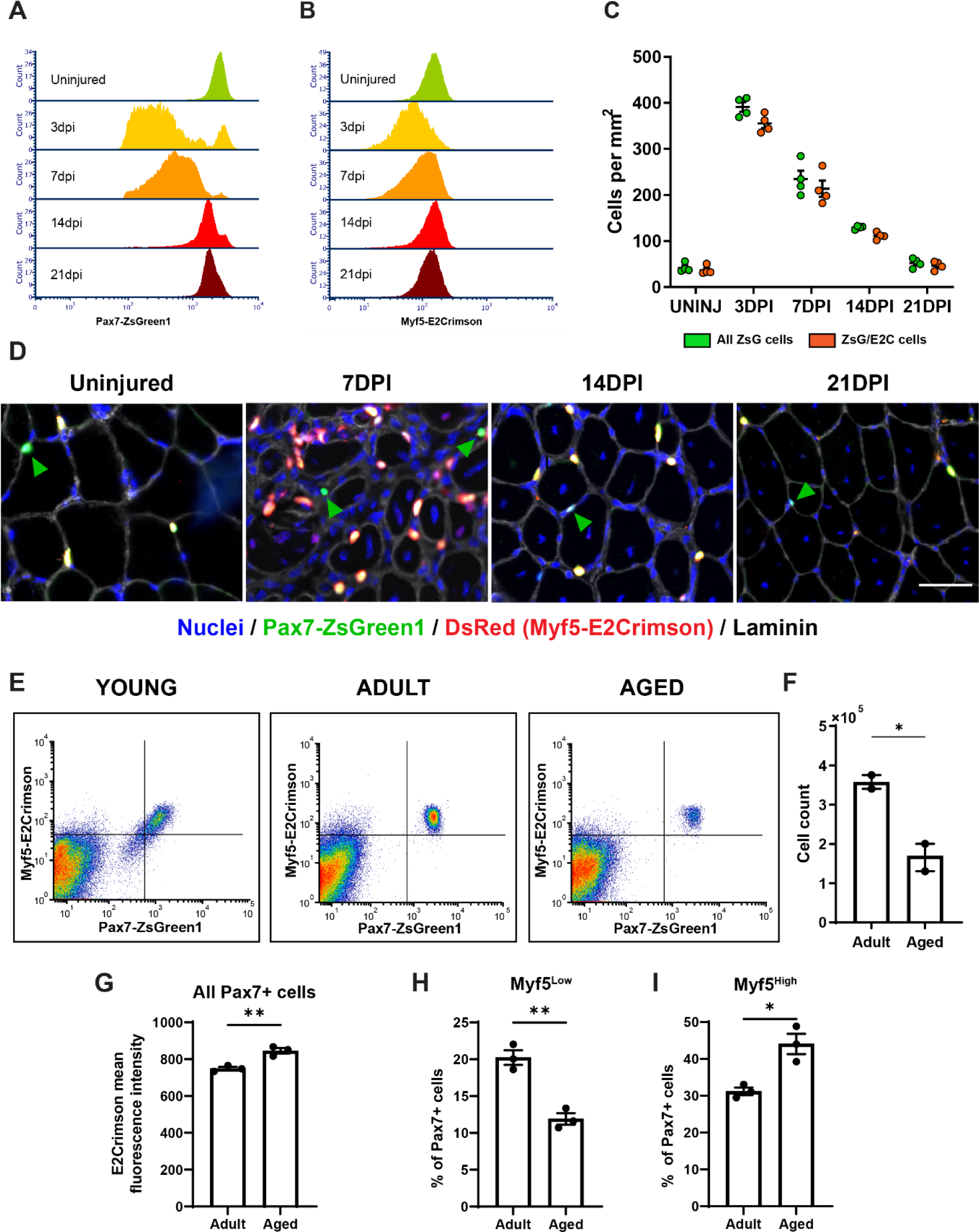
Myf5 subpopulations during adult myogenesis and aging. (**A,B**) Flow cytometry analysis of Pax7-ZsGreen1 (A) and Myf5-E2Crimson (B) fluorescence intensity dynamics in MuSCs isolated from quadriceps across under uninjured conditions and at 3 days post injury (dpi), 7dpi, 14dpi, and 21dpi. (**C,D**) Quantification (C) and representative images (D) of MuSC subpopulations from tibialis anterior histological cross-sections at multiple stages of regeneration. Scale bar = 50µm. (**E**) Representative flow cytometry profiles of MuSCs isolated from young (left), adult (middle), aged (right) StemRep mice. (**F**) Flow cytometry-based quantification of MuSC numbers in adult and aged StemRep mice. n = 4 mice, 2 pools of 2 mice. (**G**) Quantification of Myf5-E2Crimson mean fluorescence intensity in MuSCs derived from adult and aged StemRep mice. n = 3 mice per group, each datapoint represents one mouse. (**H**,**I**) Proportion of Myf5^Low^ (H) and Myf5^High^ (I) MuSCs in adult and aged StemRep mice. n = 3 mice per group, each datapoint represents one mouse. Data presented are means ± s.e.m.; *p<0.05 in a two-tailed unpaired Student’s t-test (F-I).

To study how Myf5 protein levels in MuSCs are regulated during the physiological aging process, we evaluated their fluorescence profile in hindlimb muscles of young 3 week-old (w.o.), adult 4 month-old (m.o.), and aged (24 m.o.) StemRep mice by flow cytometry (**Figures 3E,F**). Consistent with the idea that amplifying myogenic progenitors are predominant in postnatal muscle before establishing quiescence (Bachman et al., 2018; Gattazzo et al., 2020), flow cytometry analysis of 3 w.o. juvenile mice displayed fluorescence profiles for both ZsGreen1 and E2Crimson that were reminiscent of regenerating muscles (**Figures 3E** and **Figures S4A**). Compared to 4 m.o. young adult mice with a predominantly quiescent MuSC pool, aged 24 m.o. muscles contained fewer MuSCs and displayed a 13% increase in E2Crimson mean fluorescence intensity across the entire stem cell population (**Figures 3F-G**). In line with the impaired maintenance of the uncommitted MuSC pool during aging, the Myf5^High^ population became more abundant in 24 m.o. StemRep mice (**Figures 3H,I** and **Figures S4B,C**). Altogether, these data show that StemRep mice can be used to monitor the dynamics of stem cell subpopulations in different physiological contexts *in vivo*, and they support the notion that the smaller MuSC pool in aged muscles loses quiescence and becomes more committed to terminal differentiation.

### Functional heterogeneity of Myf5 subpopulations in StemRep mice

To assess the functional properties of Myf5^High^ and Myf5^Low^ MuSC subpopulations, isolated single clones were grown in high-serum conditions for ten days (**Figure 4A**). No evidence of distinct clonal efficiency was observed, suggesting that the two populations are equally potent in starting a colony. However, colonies derived from Myf5^High^ cells were on average larger (634 cells/clone) compared to the ones yielded by Myf5^Low^ clones (489 cells/clone) (**Figure 4B**). We next analyzed the distribution of clones according to their size as well as their relative frequency. This experiment revealed that 26% of Myf5^Low^ colonies generated clones with less than 250 nuclei, while only 16% of Myf5^High^ clusters fell into this category (**Figure 4C**). To check if Myf5 subpopulations differed in terms of dynamics of myogenic progression, we interrogated the changes in expression profiles of Pax7 and MyoD. By 48h and 72h post-sort, 37% and 84% of Myf5^High^ MuSCs downregulated Pax7 respectively, while 22% and 79% were negative in the Myf5^Low^ condition at the respective timepoints (**Figure 4D**). Moreover, Myf5^High^ MuSCs showed a 12% to 49% stronger induction of MyoD across the 3-day time course compared to Myf5^Low^ cells (**Figure 4E**). These results demonstrate that Myf5^Low^ MuSCs are slower to undergo myogenic commitment to differentiation than Myf5^High^ cells.

**Figure 4:**
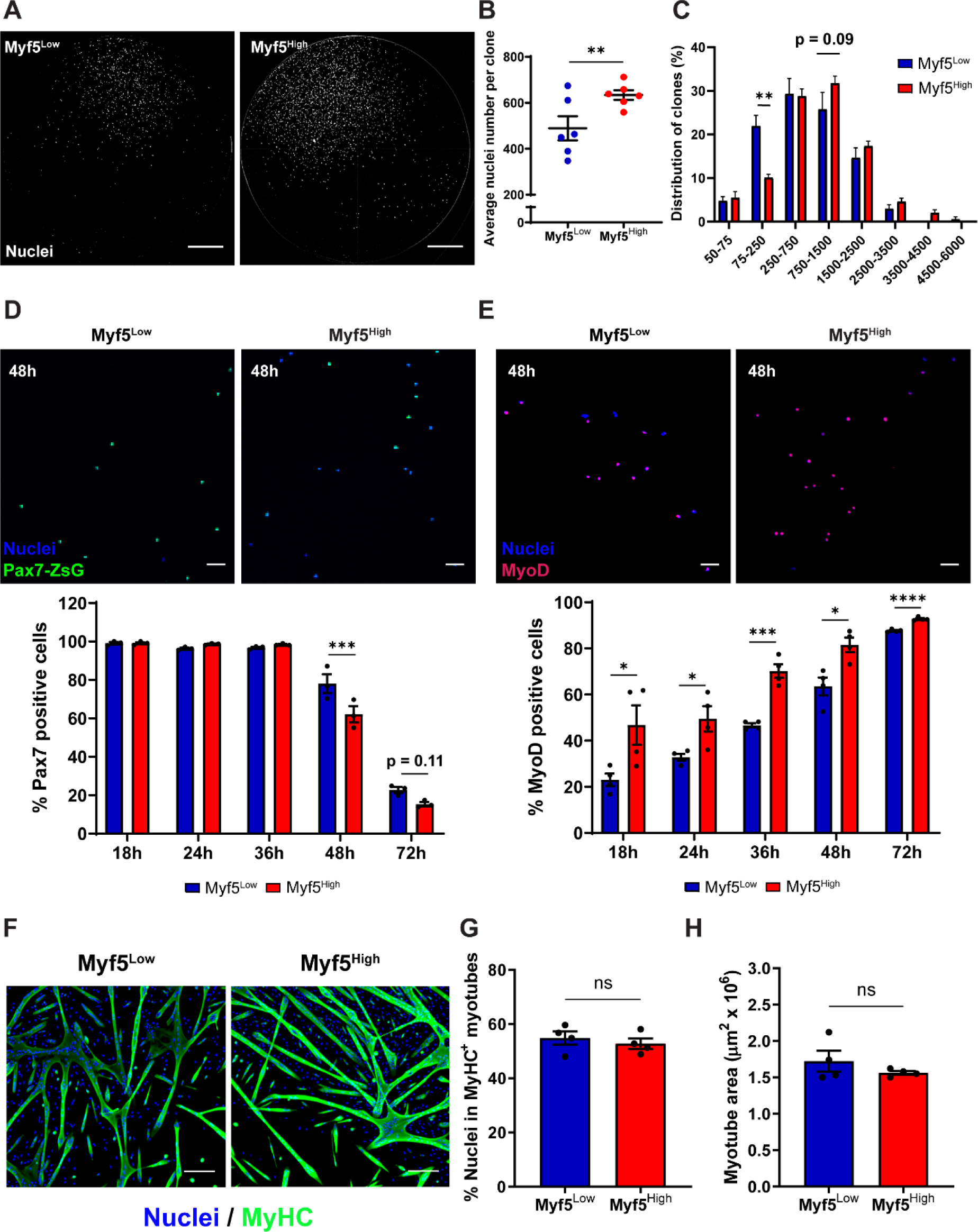
Distinct clonal and functional properties of Myf5 subpopulations. (**A-C**) Representative images (A), nuclei count (B), and size distribution (**C**) Distribution of clones from freshly isolated Myf5^High^ and Myf5^Low^ cells after 10 days of culture. n = 6 x 96-well plates from a pool of 4 mice. Scale bar = 1 mm. (**D-E**) Representative pictures (top) and quantification (bottom) of Pax7 (D) and MyoD (E) expression in Myf5^High^ and Myf5^Low^ after 48h of culture. Scale bars = 50µm. n = 4 mice. (**F-H**) Representative pictures (F) and quantification of fusion capacity (G) and myotube size (H) of Myf5 subpopulations. Myotube area was defined as the area positive for the myosin heavy chain (MyHC). Scale bar = 100µm. n = 4 mice. Data presented are means ± s.e.m.; *p<0.05, **p<0.01, ***p<0.001, ****p<0.0001 in a two-tailed unpaired Student’s t-test for each timepoint (B, C, D, E, G, H).

To assess the differentiation efficiency of the two MuSC subpopulations, Myf5^Low^ and Myf5^High^ cells were plated at confluence after isolation and medium was changed to low-serum conditions the next day (**Figure 4F**). No significant changes were observed in terms of differentiation capacity based on the number of nuclei within myotubes and their total area (**Figures 4G,H**). These observations demonstrate that the differences between the two MuSC populations are constrained to stem cell level and, once a differentiation signal is received, both cell types are equally efficient in progressing through the myogenic program.

### Myf5^Low^ MuSCs exhibit molecular and phenotypic signatures of deep quiescence

To understand how underlying molecular mechanisms translate into distinct functional properties in Myf5 subpopulations, we performed RNA-sequencing using freshly isolated Myf5^Low^ and Myf5^High^ MuSCs from adult StemRep mice. For the first time in this study, we observed sex differences in this experiment. The principal component analysis (PCA) of all expressed genes revealed a separation according to male and female mice that accounted for 30% of the total variance, while Myf5 expression accounted for 19.9% (**Figure 5A**). When examining sex-independent gene signature differences between the MuSC populations, we observed an enrichment in transcripts related to activation and differentiation in the Myf5^High^ fraction (**Figures 5B-D**). Supporting the notion that Myf5 levels control the exit from quiescence and activation, when equal numbers of freshly isolated MuSCs from StemRep mice where plated in pro-proliferative media, the nuclei count in the Myf5^Low^ condition was 21% lower at 24h, and 14% lower at 72h when compared to Myf5^High^ cells (**Figures 5E,F**). Confirming slower activation under pro-proliferative conditions, 16% fewer Myf5^Low^ MuSCs incorporated EdU at 24h when compared to the Myf5^High^ condition (**Figure 5G**). Collectively, these results demonstrate that Myf5^Low^ MuSCs in StemRep mice reside in deeper quiescence and are slower to activate than Myf5^High^ cells.

**Figure 5:**
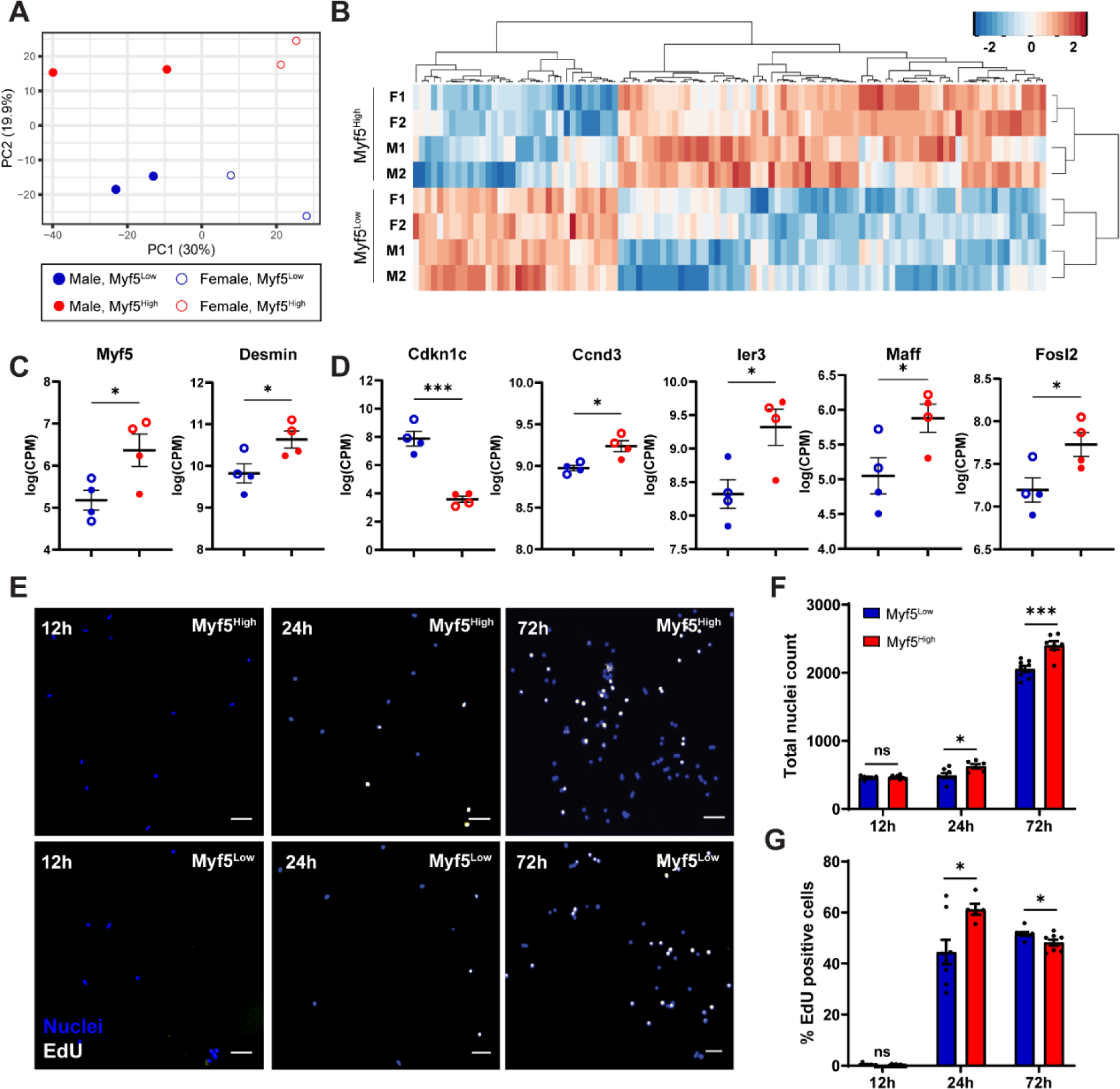
Molecular profiling of Myf5 subpopulations. (**A**) Principal component analysis of Myf5^High^ and Myf5^Low^ MuSCs isolated from male and female StemRep mice. Each datapoint represents 50,000 MuSCs isolated from a pool of 4 animals. (**B**) Heatmap of differentially expressed genes between the Myf5^High^ and Myf5^Low^ transcriptomes. (**C-D**) Differential expression of selected muscle-(C) and cell cycle-related genes (D) between Myf5 fractions in the male datasets. (**E-G**) Representative images (E) and quantification of nuclei (F) and EdU-positive cells (G) in Myf5^High^ and Myf5^Low^ subpopulations over a 3-day time course. Scale bar = 50µm. n = 4 mice. Data represent means ± s.e.m.; *p<0.05, ***p<0.001 in a two-tailed unpaired Student’s t-test for each timepoint (C, D, F, G).

Altogether, we demonstrate that StemRep mice are a versatile tool for the analysis of Pax7 and Myf5-based heterogeneity in the MuSC pool and characterize Pax7^high^Myf5^Low^ cells as an uncommitted stem cell subpopulation that resides in deep quiescence.

## Discussion

Diverse post-transcriptional mechanisms regulating mRNA stability, storage, and the rate of translation have been discovered (Shyu et al., 2008). In particular, stem cell populations that remain quiescent over much of their lifetime are able to store mRNA allowing them to be primed for efficient activation (Cheung and Rando, 2013). These considerations are particularly important with respect to the regulation of Myf5, which has been shown to be a key factor for MuSC commitment. In quiescent MuSCs, the mRNA coding for Myf5 and microRNA-31 suppressing its translation, have been shown to be stored in mRNP granules (Crist et al., 2012). Once the cells activate, mRNP granules in MuSCs dissolve and miR-31 levels are lowered leading to an accumulation of Myf5 protein. In the earliest stages of MuSC activation, this process has been suggested to be independent of transcription. Another mechanism uncoupling mRNA from protein levels in quiescent MuSCs involves translation initiation factor eIF2α (Zismanov et al., 2016). eIF2α is phosphorylated by PKR-like endoplasmic reticulum kinase (Perk) leading to a general downregulation of global translation and selectivity towards certain mRNAs. In addition, phosphorylated eIF2α is required for formation and maintenance of mRNP granules containing Myf5 and MyoD. Therefore, knock-in reporters alleles for Myf5 such as Myf5^LacZ^ or Myf5^GFP^ provide valuable insights into gene regulation, but do not always allow for an accurate prediction of protein levels (Beauchamp et al., 2000; Kassar-Duchossoy et al., 2004; Tajbakhsh et al., 1996). Depending on the targeting strategy, such alleles can also result in haploinsufficiency accompanied by a partial loss of gene product (Morrill and Amon, 2019). Thus, gene-dosage dependent effects are yet another aspect that needs to be considered when analyzing stem cell function in knock-in reporter models.

Raising antibodies to a protein of interest can help determine and study its function. However, this approach is not always successful and off-target reactivity as well as high levels of background signal are often a problem. As an alternative, epitope tagging, for instance using fluorescent proteins, is a frequently employed strategy to characterize, purify, and determine the *in vivo* localization of gene products (Brizzard, 2008). A limiting aspect of epitope tags is their potential interference with protein structure and function. The recent discovery of “self-cleaving” 2A peptides that are present in a wide range of viral families has allowed to partially bypass this problem (Liu et al., 2017). 2A-mediated self-cleavage has been suggested to involve a failure of the ribosome in making a peptide bond (Liu et al., 2017; Luke et al., 2008). Therefore, 2A peptides can be used to link a genomic locus of interest to post-translationally cleaved epitopes leaving protein function unimpaired and resulting in equimolar levels of gene product and reporter. While subcellular localization is difficult to analyze using this strategy, it is well suited to measure the amount of a given protein product across cell populations. Importantly, when quantitative assessment of a gene product based on a reporter molecule is attempted, 2A peptides have the advantage that they do not suffer from lower expression levels of downstream proteins occurring in bi-cistronic mRNA open reading frames with internal ribosomal entry site (IRES) elements (Hennecke et al., 2001). Next to P2A derived from porcine teschovirus-1 2A, the T2A peptide derived from Thosea Asigna Virus 2A, has been shown to have particularly high cleavage efficiency in mammalian cells (Donnelly et al., 2001; Kim et al., 2011; Szymczak et al., 2004). Here, we used T2A to link the last exon of the Pax7 and Myf5 genes to two different nuclear targeted fluorescent proteins, allowing for assessment of the real-time dynamics of these two factors on the protein level *in vivo* and after isolation *in vitro*. Nuclear localization signals (NLS) are short peptides that mediate the translocation of proteins from the cytoplasm into the nucleus (Mahato et al., 1999). MuSC numbers and function are often analyzed using thin skeletal muscle cross-sections. Due to their two-dimensional nature, it is challenging to quantify the confines of cells marked by cytoplasmatic or membrane proteins in this type of sample preparation. The addition of NLS to the reporter proteins in StemRep mice has solved this problem and allows for convenient identification, quantification, and isolation of MuSCs with different levels of Pax7 and Myf5. Thus, the StemRep model will facilitate the study of MuSC heterogeneity under different physiological and pathological conditions.

To demonstrate a specific application of StemRep mice in studying physiological changes, we examined the influence of age on the Myf5^Low^ and Myf5^High^ MuSCs. When compared to adult 4 m.o. StemRep mice, MuSCs in young 3 w.o. animals showed a wider range of fluorescence intensities akin to regenerating muscles. This correlates well with the observation that MuSCs in postnatal muscles are still highly active and contribute to the growth of myofibers (Bachman and Chakkalakal, 2022). In contrast aging is well known to lead to both a loss of MuSC numbers and function (Mashinchian et al., 2018). We observed that aged 24 m.o. StemRep mice showed higher levels of Myf5 across the entire MuSC population. This result supports the notion that aging causes the break of MuSC quiescence which impairs maintenance of the uncommitted stem cell pool and ultimately leads to regenerative failure (Ancel et al., 2021; Brack et al., 2007). Next to aging, an impaired healing capacity of skeletal muscle has been described to accompany many other conditions, including muscular dystrophy and cancer cachexia (Deprez et al., 2023; Mashinchian et al., 2018). Therefore, future experiments using the StemRep alleles on disease backgrounds may provide important novel insights into pathogenesis and might unravel therapeutic approaches to target specific cell MuSC subpopulations.

We observed that all cells expressing the canonical MuSC cell surface markers α7-integrin and CD34 were positive for Pax7-ZsGreen1 fluorescence. Live-cell imaging, muscle regeneration after experimental injury, and flow cytometry analysis at different ages confirmed that Pax7-ZsGreen1 and Myf5-E2Crimson in StemRep mice are dynamically regulated and faithfully recapitulate the temporal endogenous expression patterns of Pax7 and Myf5 (Bentzinger et al., 2012). About 10% of Myf5^Low^ MuSCs in StemRep mice are present in adult homeostatic muscle and persist throughout regenerative myogenesis. This result suggests that this population corresponds to the Myf5-lineage negative satellite stem cell MuSC population that was described in Myf5-Cre;ROSA26-YFP mice (Kuang et al., 2007). Notably, we found that Myf5^Low^ cells contain significantly increased levels of Pax7 protein, which uncovers an interesting parallel between MuSC populations defined by Myf5 levels and the Pax7-nGFP^Hi^ subpopulation observed in mice with a nuclear GFP inserted into the first exon of Pax7 gene that were shown by Rocheteau et al. to undergo asymmetric DNA strand segregation (Rocheteau et al., 2012). For the first time, our results demonstrate that the Pax7^High^ and Myf5^Low^ MuSC subpopulations likely correspond to the same fraction of cells.

Although both Myf5^Low^ and Myf5^High^ MuSCs isolated from StemRep mice were equally capable of forming stem cell colonies and myotubes, Myf5^Low^ cells were slower to enter cell cycle, proliferate, and commit to myogenic differentiation, demonstrating that this subpopulation resides in a deeper state of quiescence compared to Myf5^High^ cells. This observation was supported by transcriptomic profiling of the Myf5 subpopulations, showing that Myf5^High^ cells are enriched in transcripts implicated in early activation. The discovery that Myf5^Low^ MuSCs are residing in deep quiescence is yet another common denominator with previous studies describing superior stemness characteristics of the most dormant stem cell populations in skeletal muscle (García-Prat et al., 2020; Rocheteau et al., 2012; Rodgers et al., 2014). Thus, we present the unifying hypothesis that the adult MuSC pool contains a subpopulation of stem cells in deep quiescence characterized by high levels of Pax7 and low levels of myogenic commitment factors such as Myf5 and MyoD. This population of cells is likely the least committed fraction of MuSCs present in adult skeletal muscle and is responsible for the maintenance of the stem cell pool and, ultimately, regenerative capacity.

Our study draws several important parallels with previous work, fills in long-standing gaps in our understanding of MuSC subpopulations, and establishes the StemRep line as a novel reagent for the interrogation of stem cell heterogeneity in skeletal muscle. Our results unify multiple existing theories about the quintessential stem cell population in skeletal muscle and point towards a fraction of roughly 10% of MuSCs that are metabolically inactive, reside in deep quiescence, and are marked by high Pax7 and low Myf5 protein levels. The broad availability of StemRep mice to the field will enable future in-depth studies addressing the molecular characteristics of this “high-stemness” MuSC population in health and disease.

## Acknowledgements

We thank the NIHS community for critical discussions of the results and manuscript. C.F.B. is supported by the Canadian Institutes of Health Research (CIHR, PJT-162442), the Natural Sciences and Engineering Research Council of Canada (NSERC, RGPIN-2017-05490), the Fonds de Recherche du Québec - Santé (FRQS, Dossiers 296357 and 34813), and the Fonds de Recherche du Québec - Nature et Technologies (FRQNT, Dossier 331297).

## Author contributions

S.A., P.S., and J.N.F. designed the experimental strategy, interpreted the results, and wrote the manuscript. P.S. and J.N.F. led the project. C.F.B. ideated and conceived the mouse line and edited the manuscript. S.A., J.M., F.S., S.K. and P.S. performed experiments and analyzed data. S.A., F.S., L.T., M.O. and G.D. designed the flow cytometry strategy, performed the sorts and analyzed the results. J.L.S.G. supported animal work. E.M. analyzed transcriptomic experiments. G.J. and S.M. supported imaging and genomics. All authors reviewed the manuscript.

## Experimental Procedures

### Data and materials availability

The newly generated StemRep mice are being submitted to the Jackson Labs mouse repository (https://mice.jax.org/) and will be freely available to the academic community upon publication of this manuscript. All software used were freely or commercially available. Any additional information required to reanalyze the data reported in this paper is available upon justified request to the corresponding authors. RNA sequencing data have been deposited to GEO under accession number GSE207321. Other materials are available upon justified request within the limits of non-renewable materials.

### Generation of the Pax7/Myf5StemRep reporter mice

The Pax7-ZsGreen1 (C57BL/6NTac-Pax7tm3955(T2A-ZsGreen)Tac) and Myf5-E2Crimson (C57BL/6NTac-Myf5tm3956(T2A-E2Crimson)Tac) knock-in mice were generated by Taconic Biosciences via a specific service contract designed by lead authors. The targeting strategy was based on NCBI transcripts NM_011039.2 (Pax7) and NM_008656.5 (Myf5). For the Pax7-ZsGreen1 line, the targeting vector contained the *Pax7* open reading frame (ORF) together with a Kozak sequence to allow protein translation initiation. The DNA sequences of the T2A self-cleaving peptide and the NLS-ZsGreen1 ORF were inserted between the last amino acid and the translation termination codon in exon 9. For the Myf5-E2Crimson line, the T2A and the NLS-E2Crimson ORF were inserted between the last amino acid and the translation termination codon in exon 3. A Puromycin resistance cassette flanked by FRT sites was inserted into intron 8 of the *Pax7* gene and downstream of the 3’ untranslated region of the *Myf5* gene. The targeting vectors were generated using BAC clones from the C57BL/6J RPCI-23 BAC library which were then transfected into the Taconic Biosciences C57BL/6N Tac ES cell line. Homologous recombinant clones were isolated using positive Puromycin and negative Thymidine kinase selection. Final knock-in alleles were generated by in-vivo Flp-mediated removal of the selection marker. The resulting transgenics were intercrossed to generate double heterozygotes, which were used to start generating the double homozygous StemRep experimental animals used in this study.

### Animal housing and ethics

All *in vivo* experiments and protocols were performed in compliance with the regulations of the Swiss Animal Experimentation Ordinance and approved by the internal ethics committee of Nestlé Research and the ethical committee of the canton de Vaud under license VD3331. Mice were housed under standard conditions (up to 5 mice per cage) and allowed access to food and water *ad libitum*. Unless otherwise specified, all experiments were conducted using both males and females at an approx. 1:1 ratio. All data was systematically analyzed for sex differences and the results were reported if divergence was observed. For aging studies, animals were bred and housed in the same facility using housing conditions described above.

### Genotyping

To identify animals carrying the Pax7-ZsGreen1 and Myf5-E2Crimson constructs, genomic DNA was isolated using the KAPA Express Extraction protocol (KAPA Biosystems, # KB7101). Briefly, ear snips were lysed in 1X KAPA Express Extract Buffer and 2U of KAPA Express Extract Enzyme diluted in PCR grade water for 10 min at 75°C and 5 min at 95°C. DNA fragments were then amplified in PCR buffer composed of 2 mM MgCl2, 0.2 mM dNTPs (Roche, #07958846001), 25 µM of forward and reverse primers, 1X High Fidelity Buffer and 1U Platinum Taq DNA Polymerase High Fidelity (Thermo Fisher Scientific, #11304011) in PCR grade water. The following primers were used: Pax7-ZsG_For: 5’-AGA-TTGGCAGGTGTGTAAAGG-3’, Pax7-ZsG_Rev: 5’-TTTATCCATCTTCTACTCCATCCC-3’, and Myf5-E2C_For: 5’-GCATCTGTGAGATGGATGGG-3’, Myf5-E2C_Rev: 5’-AGGACTACACAGCAAAACCTGTG-3’. PCR products were generated under the following conditions: 95°C hold for 5 min, 35 cycles of 95°C for 30 sec, 60°C for 30 sec and 72°C of 1 min, followed by a 72°C hold for 10 min. Pax7-ZsGreen1 and Myf5-E2Crimson constructs yield fragments of 356 and 345 base pairs, respectively.

### Isolation of Myf5 subpopulations by FACS

MuSCs from StemRep mice were isolated as previously described (Lukjanenko et al., 2016). Hindlimb muscles from injured or uninjured mice were collected, minced, and digested with Dispase II (2.5 U/ml) (Roche), Collagenase B (0.2%) (Roche) and MgCl2 (5 mM) at 37°C. The preparation was then filtered sequentially through 100 micron and 30 micron filters. For initial validation of the Pax7/Myf5 model, cells were incubated at 4°C for 30 min with antibodies against CD45 (Invitrogen, #MCD4528, 1/25), CD31 (Invitrogen, #RM5228, 1/25), CD11b (Invitrogen, #RM2828, 1/25), CD34 (BD Biosciences, #560238, 1/60), Ly-6A/E (BD Biosciences, #561021, 1/150) and α7-integrin (R&D Systems, #FAB3518N, 1/30). Myf5 subpopulations identified as Pax7-ZsGreen1 positive and Myf5-E2Crimson high or low were isolated using a Beckman Coulter Astrios Cell sorter.

### Live-cell imaging

MuSCs were freshly isolated and seeded on 0.2% gelatin-coated 96 well plates at a density of 2,000 cells per well in growing conditions and incubated for 4 days in an IncuCyte SX5 Live-Cell Analysis System (Sartorius) at 37°C and 5% CO2. Cells were imaged every 12 h and cell growth and green fluorescence metrics were extracted and analyzed using the IncuCyte ZOOM software (Sartorius).

### Muscle regeneration

Muscle regeneration was induced by intramuscular injection of 25 µl, 50 µl, and 50 µl of 20 µM cardiotoxin (CTX, Latoxan) into the *Tibialis anterior* (TA), *Gastrocnemius* (GC), and *Quadriceps femoris* (QD) muscles, respectively, under a short anesthesia using 2% isoflurane. For analgesia, buprenorphine was administered at a dosage of 0.1 mg/kg subcutaneously before the intramuscular injection followed by a second dose 24 h after. Mice were sacrificed by CO2 exposure 3-, 7-, 14- or 21- days post-injury and muscles were collected for analysis. TA muscles were mounted on 6% tragacanth gum (Sigma, #G1128) and frozen in isopentane (Sigma, #M32631) cooled with liquid nitrogen for histological analysis. GC and QD muscles were immediately processed after harvest for analysis by flow cytometry.

### Histology and image analysis

Frozen TA muscles were sectioned at 10 µm with a cryostat (Leica Biosystems). Samples were fixed with 4% PFA (EMS, #157-4-100), and permeabilized in 0.5% Triton X-100 (Sigma, X100) diluted in PBS (PBTX). Sections were further blocked in 4% BSA for 2 h. For immunostaining the slides were incubated with primary antibodies anti-DsRed (Takara Bio, #632496, 1/1000), anti-Pax7 (purified hybridoma, DSHB, 1/1000), and anti-laminin (LSBio, # LS-C96142, 1/1000). Nuclei were labelled with Hoechst 33342 (Sigma, #B2261) during secondary antibody incubation (Thermo Fisher Scientific, #A21245 and #A21437). Slides were then mounted using Dako fluorescent mounting medium (Agilent, #S302380-2) and imaged with an Olympus VS120 slide scanner. Images were analyzed using the VS-ASW FL software measurement tool. The number of Pax7 and Myf5 positive cells was determined by manually counting injured areas on muscle sections.

### MuSC-derived myoblasts and staining

FACS-isolated Myf5^High^ and Myf5^Low^ cells were directly plated onto 0.2% gelatin-coated 384-well plates in growth medium (DMEM supplemented with 20% heat-inactivated FBS, 10% horse serum, 2.5 ng/ml bFGF, 1% P/S and 1% L-Glutamine). To assess proliferation, 5μM EdU was added in the medium for 2 h prior to the assessed timepoints. EdU incorporation was revealed using the Click-iT assay (Thermo Fisher Scientific, #C10338) according to manufacturer’s instructions. Briefly, cells were fixed during 15 min in 4% PFA, permeabilized during 20 min in PBTX 0.5%, stained with the Click-iT reaction mix and counterstained with Hoechst 33342 (Sigma, #B2261). To assess differentiation, 5,000 Myf5^High^ and Myf5^Low^ cells were seeded onto 0.2% gelatin-coated 384-well plates in growing medium. 12 h post-plating, cells were switched to differentiating conditions (DMEM with 5% horse serum and 1% P/S) and fixed after 4 days. Immunostaining was performed by blocking for 1h in 4% BSA (Sigma, #A8022) and staining with anti-DsRed (Takara Bio, #632496, 1/1000), anti-Pax7 (purified hybridoma, DSHB, 1/1000), anti-MyoD (LsBio, # LS-C143580-100, 1/500), anti-Myogenin (Abcam, #124800, 1/500), or anti-MyHC (Merck, #05-716, 1/200) and then with appropriate secondary antibodies (Thermo Fisher Scientific, #A21245 and #A32727) and Hoechst 33342 (Sigma, #B2261). Images were acquired using the ImageXpress (Molecular Devices) platform and quantifications were performed using the MetaXpress software (Molecular Devices). Exclusion of staining artifacts was performed based on morphological analysis and myotubes were detected based on the segmentation of MyHC staining. Multi-wavelength cell scoring was performed by identifying positive and negative populations based on a minimum intensity threshold for each marker using an automated image processing module.

### RNA-sequencing of Myf5^High^ and Myf5^Low^ subpopulations

Myf5^High^ and Myf5^Low^ MuSCs were isolated from a pool of 4 male and 4 female mice and repeated in two independent experiments and total RNA was extracted using Agencourt RNAdvance tissue Kit (Beckman Coulter, #A32646) with a quality score of RQN 9.3 to 10. Libraries were constructed from cDNA generated and amplified (21 Cycles) from 20 ng of RNA, following the user guide with QuantSeq 3’ mRNA-Seq Library Prep Kit FWD for Illumina (Lexogen) without any modification. Libraries were quantified with Quant it Picogreen (Invitrogen, #10545213). The library sizes were controlled with the High Sensitivity NGS Fragment Analysis kit on a Fragment Analyzer (Agilent). Sequencing was performed on HiSeq 2500 with Rapid V2 chemistry SR 65 cycles (Illumina) using two different flow cells loaded at 2pM and with 3% Phix. Primary data QC was performed during the sequencing run to ensure the optimal flow cell loading (cluster density) and check the quality metrics of the sequencing run.

### Statistical analyses

Unless otherwise stated, data were analyzed using the Prism 9 software package and represented as mean ± SEM. Readouts used in this study were assessed for normality based on the distribution of historical values across multiple experiments using a D’Agostino-Pearson test. Two conditions comparisons were analyzed using a student’s t-test.

### RNA-sequencing statistics

Sequencing data were demultiplexed with Bcl2FastQ and transformed into fastq files using casava v1.8.2. The fastq files were then aligned against the reference genome GRCm38 using RNAstar v2.5.3a (Dobin et al., 2013). Counts per gene from the bam files were generated using htseq_count v2.16.2 (Anders et al., 2015). All transcriptomic analyses were performed using R (version 3.6.1) and relevant Bioconductor packages. After selecting genes with more than 8 counts per million in at least 4 samples, RNA-sequencing data were normalized by the trimmed mean of M-values (TMM) method as implemented in function calcNormFactors in edgeR (Robinson et al., 2010). Differentially expressed genes were defined by fitting a quasi-likelihood negative binomial generalized log-linear model to count data using glmQLFTest function in edgeR with mouse sex and Myf5 expression as explanatory variables.

## Supplementary Figures

**Figure S1 (related to Figure 1):**
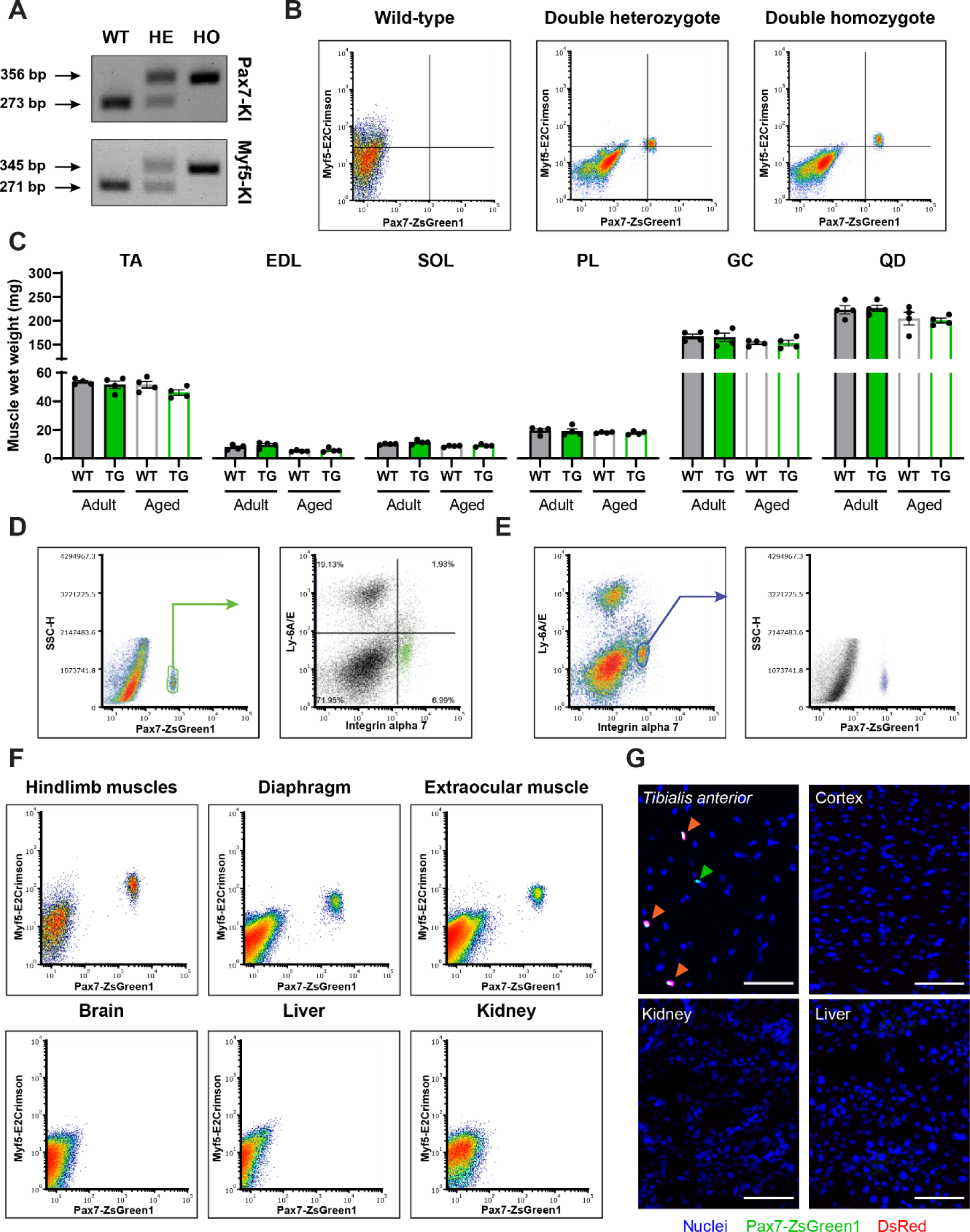
Generation and validation of StemRep mice. (**A**) PCR genotyping of WT, heterozygous, and homozygous mice for Pax7-ZsGreen1 (top) and Myf5-E2Crimson (bottom) knock-in. (**B**) Representative flow cytometry plots of ZsGreen1 and E2Crimson fluorescence in MuSCs isolated from skeletal muscle (hindlimb) of double heterozygous (left) and double heterozygous (right) StemRep mice. (**C**) Analysis of WT and transgenic hindlimb muscle weights from young and aged mice. TA, *tibialis anterior*; EDL, *extensor digitorum longus*; SOL, *soleus*; PL, *plantaris*; GC, *gastrocnemius*; and QD, *quadriceps femoris*. Data presented are means ± s.e.m. (**D,E**) Representative flow cytometry plots of ZsGreen1 fluorescence in the CD11b-/CD31-/CD45-/Sca1-/CD34+/Itga7+ MuSC population from StemRep mice (D) and of the antigen signature in ZsGreen1 cells (E). (**F,G**) Representative flow cytometry profiles (F) and pictures (G) of Pax7-ZsGreen1 and Myf5-E2Crimson fluorescence in cross sections of skeletal muscles, brain, liver, and kidney. My5^High^ MuSCs are indicated with an orange arrowhead while green arrowheads designate Myf5^Low^ cells. Scale bar = 100µm.

**Figure S2 (related to Figure 1):**
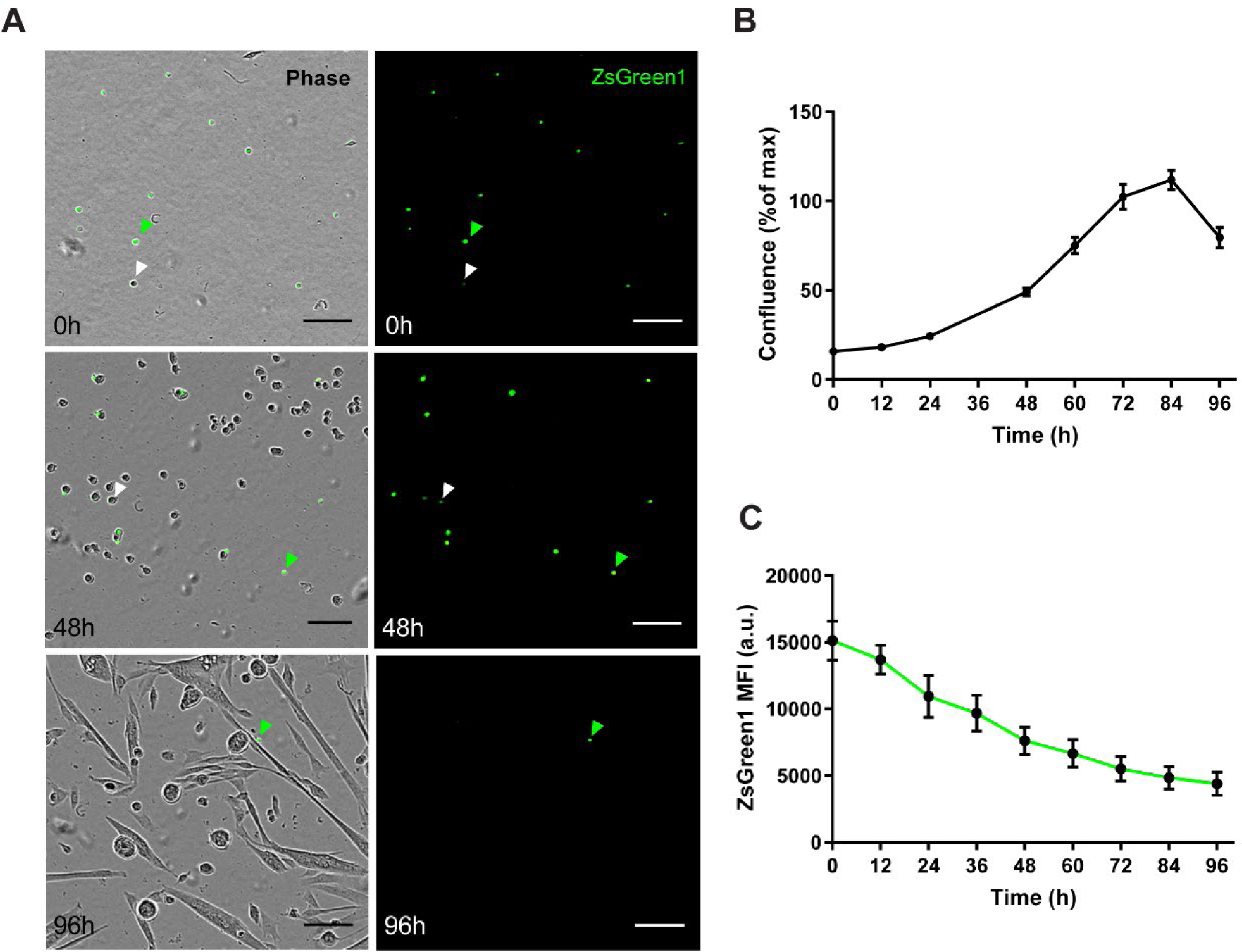
Temporal dynamics of Pax7-ZsGreen1 cells. (**A-C**) Representative pictures (A) and quantification of cell growth (B) and ZsGreen1 mean fluorescence intensity of MuSCs isolated from StemRep mice (C) by live-cell imaging over a time course of 4 days. Scale bar = 100µm. Green arrowheads denote strongly zsGreen1 positive cells, white arrowheads denote cells with low zsGreen1 levels.

**Figure S3 (related to Figure 2):**
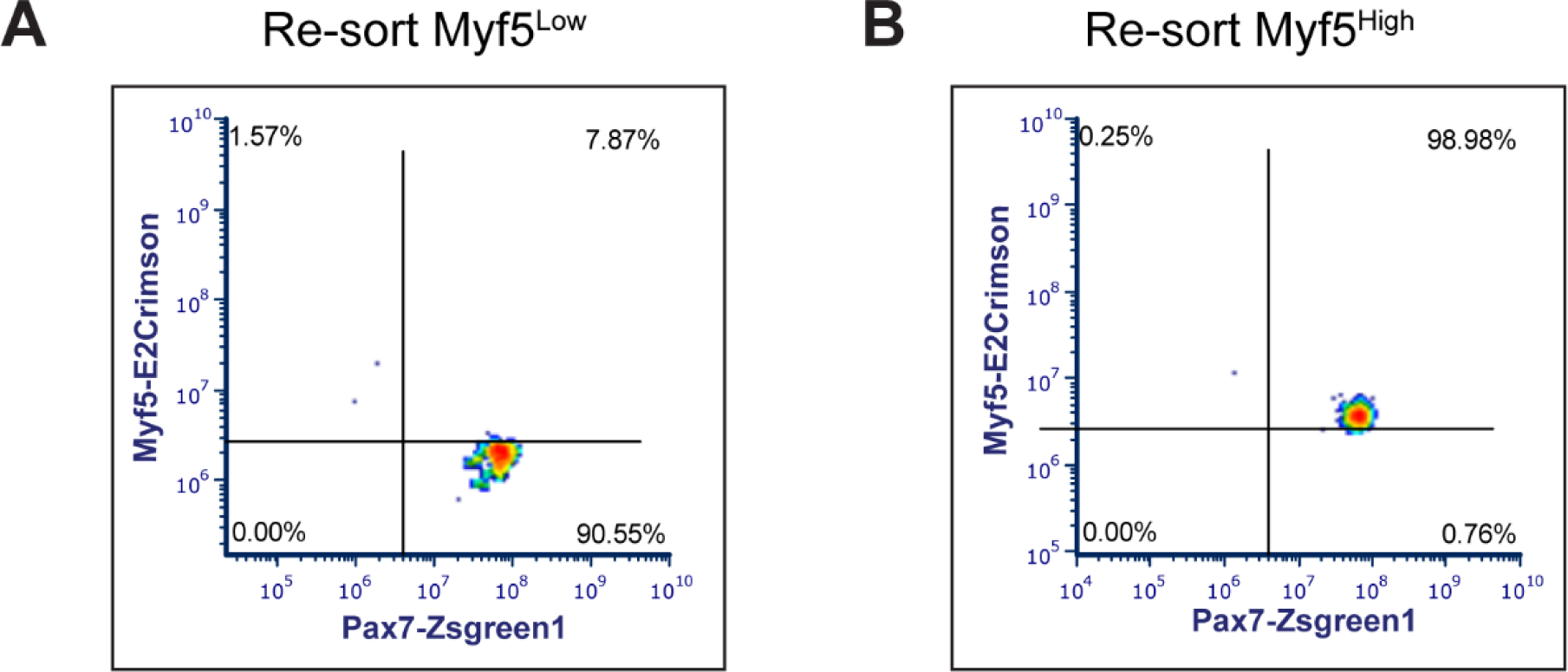
Flow cytometry analysis of Myf5 subpopulations. (**A,B**) Representative flow cytometry profiles of Myf5^Low^ (A) and Myf5^High^ (B) cells re-sorted following isolation.

**Figure S4 (related to Figure 3):**
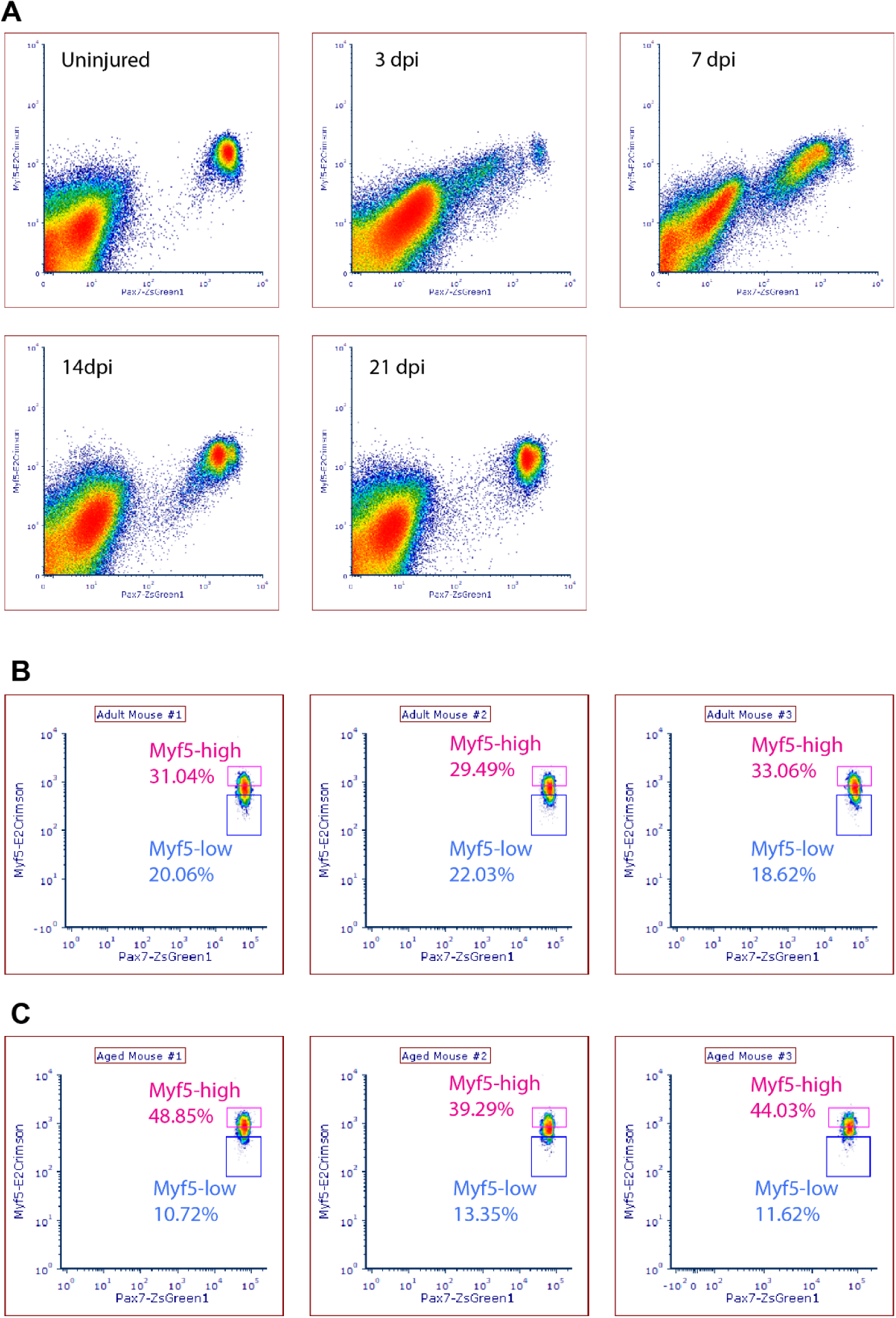
Flow cytometry analysis of Myf5 subpopulations in regenerating and aged skeletal muscle. (**A**) Representative flow cytometry profiles of Pax7-ZsGreen1 and Myf5-E2Crimson fluorescence in uninjured hindlimb muscles and in GC and QD muscles 3, 7, 14, and 21 dpi. n = pool of 4 mice per group. (**B**,**C**) Representative flow cytometry profiles of Pax7-ZsGreen1 and Myf5-E2Crimson fluorescence in adult (B), and aged (C). n = 3 mice per group, each plot represents the profile of uninjured hindlimb muscles from one mouse.

## References

Ancel, S., Stuelsatz, P., and Feige, J.N. (2021). Muscle Stem Cell Quiescence: Controlling Stemness by Staying Asleep. Trends Cell Biol 31, 556–568. https://doi.org/10.1016/j.tcb.2021.02.006.

Anders, S., Pyl, P.T., and Huber, W. (2015). HTSeq--a Python framework to work with high-throughput sequencing data. Bioinformatics 31, 166–169. https://doi.org/10.1093/bioinformatics/btu638.

Bachman, J.F., and Chakkalakal, J.V. (2022). Insights into muscle stem cell dynamics during postnatal development. FEBS J 289, 2710–2722. https://doi.org/10.1111/febs.15856.

Bachman, J.F., Klose, A., Liu, W., Paris, N.D., Blanc, R.S., Schmalz, M., Knapp, E., and Chakkalakal, J.V. (2018). Prepubertal skeletal muscle growth requires Pax7-expressing satellite cell-derived myonuclear contribution. Development 145. https://doi.org/10.1242/dev.167197.

Beauchamp, J.R., Heslop, L., Yu, D.S., Tajbakhsh, S., Kelly, R.G., Wernig, A., Buckingham, M.E., Partridge, T.A., and Zammit, P.S. (2000). Expression of CD34 and Myf5 defines the majority of quiescent adult skeletal muscle satellite cells. J Cell Biol 151, 1221–1234. https://doi.org/10.1083/jcb.151.6.1221.

Bentzinger, C.F., Wang, Y.X., and Rudnicki, M.A. (2012). Building muscle: molecular regulation of myogenesis. Cold Spring Harb Perspect Biol 4. https://doi.org/10.1101/cshperspect.a008342.

Brack, A.S., Conboy, M.J., Roy, S., Lee, M., Kuo, C.J., Keller, C., and Rando, T.A. (2007). Increased Wnt signaling during aging alters muscle stem cell fate and increases fibrosis. Science 317, 807–810. https://doi.org/10.1126/science.1144090.

Brizzard, B. (2008). Epitope tagging. Biotechniques 44, 693–695. https://doi.org/10.2144/000112841.

Cheung, T.H., and Rando, T.A. (2013). Molecular regulation of stem cell quiescence. Nat Rev Mol Cell Biol 14, 329–340. https://doi.org/10.1038/nrm3591.

Crist, C.G., Montarras, D., and Buckingham, M. (2012). Muscle satellite cells are primed for myogenesis but maintain quiescence with sequestration of Myf5 mRNA targeted by microRNA-31 in mRNP granules. Cell Stem Cell 11, 118–126. https://doi.org/10.1016/j.stem.2012.03.011.

Deprez, A., Orfi, Z., Rieger, L., and Dumont, N.A. (2023). Impaired muscle stem cell function and abnormal myogenesis in acquired myopathies. Biosci Rep 43, BSR20220284. https://doi.org/10.1042/BSR20220284.

Der Vartanian, A., Quétin, M., Michineau, S., Auradé, F., Hayashi, S., Dubois, C., Rocancourt, D., Drayton-Libotte, B., Szegedi, A., Buckingham, M., et al. (2019). PAX3 Confers Functional Heterogeneity in Skeletal Muscle Stem Cell Responses to Environmental Stress. Cell Stem Cell 24, 958–973.e9. https://doi.org/10.1016/j.stem.2019.03.019.

Dobin, A., Davis, C.A., Schlesinger, F., Drenkow, J., Zaleski, C., Jha, S., Batut, P., Chaisson, M., and Gingeras, T.R. (2013). STAR: ultrafast universal RNA-seq aligner. Bioinformatics 29, 15–21. https://doi.org/10.1093/bioinformatics/bts635.

Donnelly, M.L.L., Hughes, L.E., Luke, G., Mendoza, H., Ten Dam, E., Gani, D., and Ryan, M.D. (2001). The “cleavage” activities of foot-and-mouth disease virus 2A site-directed mutants and naturally occurring “2A-like” sequences. J Gen Virol 82, 1027–1041. https://doi.org/10.1099/0022-1317-82-5-1027.

Dumont, N.A., Wang, Y.X., and Rudnicki, M.A. (2015). Intrinsic and extrinsic mechanisms regulating satellite cell function. Development 142, 1572–1581. https://doi.org/10.1242/dev.114223.

Englund, D.A., Murach, K.A., Dungan, C.M., Figueiredo, V.C., Vechetti, I.J., Dupont-Versteegden, E.E., McCarthy, J.J., and Peterson, C.A. (2020). Depletion of resident muscle stem cells negatively impacts running volume, physical function, and muscle fiber hypertrophy in response to lifelong physical activity. Am J Physiol Cell Physiol 318, C1178–c1188. https://doi.org/10.1152/ajpcell.00090.2020.

García-Prat, L., Perdiguero, E., Alonso-Martín, S., Dell’Orso, S., Ravichandran, S., Brooks, S.R., Juan, A.H., Campanario, S., Jiang, K., Hong, X., et al. (2020). FoxO maintains a genuine muscle stem-cell quiescent state until geriatric age. Nat Cell Biol 22, 1307–1318. https://doi.org/10.1038/s41556-020-00593-7.

Gattazzo, F., Laurent, B., Relaix, F., Rouard, H., and Didier, N. (2020). Distinct Phases of Postnatal Skeletal Muscle Growth Govern the Progressive Establishment of Muscle Stem Cell Quiescence. Stem Cell Reports 15, 597–611. https://doi.org/10.1016/j.stemcr.2020.07.011.

Gayraud-Morel, B., Chrétien, F., Jory, A., Sambasivan, R., Negroni, E., Flamant, P., Soubigou, G., Coppée, J.Y., Di Santo, J., Cumano, A., et al. (2012). Myf5 haploinsufficiency reveals distinct cell fate potentials for adult skeletal muscle stem cells. J Cell Sci 125, 1738–1749. https://doi.org/10.1242/jcs.097006.

Hennecke, M., Kwissa, M., Metzger, K., Oumard, A., Kröger, A., Schirmbeck, R., Reimann, J., and Hauser, H. (2001). Composition and arrangement of genes define the strength of IRES-driven translation in bicistronic mRNAs. Nucleic Acids Res 29, 3327–3334. https://doi.org/10.1093/nar/29.16.3327.

Kassar-Duchossoy, L., Gayraud-Morel, B., Gomès, D., Rocancourt, D., Buckingham, M., Shinin, V., and Tajbakhsh, S. (2004). Mrf4 determines skeletal muscle identity in Myf5:Myod double-mutant mice. Nature 431, 466–471. https://doi.org/10.1038/nature02876.

Kim, J.H., Lee, S.-R., Li, L.-H., Park, H.-J., Park, J.-H., Lee, K.Y., Kim, M.-K., Shin, B.A., and Choi, S.-Y. (2011). High cleavage efficiency of a 2A peptide derived from porcine teschovirus-1 in human cell lines, zebrafish and mice. PLoS One 6, e18556. https://doi.org/10.1371/journal.pone.0018556.

Kuang, S., Kuroda, K., Le Grand, F., and Rudnicki, M.A. (2007). Asymmetric self-renewal and commitment of satellite stem cells in muscle. Cell 129, 999–1010. https://doi.org/10.1016/j.cell.2007.03.044.

Lepper, C., Partridge, T.A., and Fan, C.M. (2011). An absolute requirement for Pax7-positive satellite cells in acute injury-induced skeletal muscle regeneration. Development 138, 3639–3646. https://doi.org/10.1242/dev.067595.

Liu, Z., Chen, O., Wall, J.B.J., Zheng, M., Zhou, Y., Wang, L., Vaseghi, H.R., Qian, L., and Liu, J. (2017). Systematic comparison of 2A peptides for cloning multi-genes in a polycistronic vector. Sci Rep 7, 2193. https://doi.org/10.1038/s41598-017-02460-2.

Luke, G.A., de Felipe, P., Lukashev, A., Kallioinen, S.E., Bruno, E.A., and Ryan, M.D. (2008). Occurrence, function and evolutionary origins of “2A-like” sequences in virus genomes. J Gen Virol 89, 1036–1042. https://doi.org/10.1099/vir.0.83428-0.

Lukjanenko, L., Jung, M.J., Hegde, N., Perruisseau-Carrier, C., Migliavacca, E., Rozo, M., Karaz, S., Jacot, G., Schmidt, M., Li, L., et al. (2016). Loss of fibronectin from the aged stem cell niche affects the regenerative capacity of skeletal muscle in mice. Nat Med 22, 897–905. https://doi.org/10.1038/nm.4126.

Maesner, C.C., Almada, A.E., and Wagers, A.J. (2016). Established cell surface markers efficiently isolate highly overlapping populations of skeletal muscle satellite cells by fluorescence-activated cell sorting. Skelet Muscle 6, 35. https://doi.org/10.1186/s13395-016-0106-6.

Mahato, R.I., Smith, L.C., and Rolland, A. (1999). 4 - Pharmaceutical Perspectives of Nonviral Gene Therapy. In Advances in Genetics, J.C. Hall, J.C. Dunlap, T. Friedmann, and F. Giannelli, eds. (Academic Press), pp. 95–156.

Mashinchian, O., Pisconti, A., Le Moal, E., and Bentzinger, C.F. (2018). The Muscle Stem Cell Niche in Health and Disease. Curr Top Dev Biol 126, 23–65. https://doi.org/10.1016/bs.ctdb.2017.08.003.

Matz, M.V., Fradkov, A.F., Labas, Y.A., Savitsky, A.P., Zaraisky, A.G., Markelov, M.L., and Lukyanov, S.A. (1999). Fluorescent proteins from nonbioluminescent Anthozoa species. Nat Biotechnol 17, 969–973. https://doi.org/10.1038/13657.

Mauro, A. (1961). Satellite cell of skeletal muscle fibers. J Biophys Biochem Cytol 9, 493–495. https://doi.org/10.1083/jcb.9.2.493.

Morrill, S.A., and Amon, A. (2019). Why haploinsufficiency persists. Proc Natl Acad Sci U S A 116, 11866–11871. https://doi.org/10.1073/pnas.1900437116.

Olguin, H.C., and Olwin, B.B. (2004). Pax-7 up-regulation inhibits myogenesis and cell cycle progression in satellite cells: a potential mechanism for self-renewal. Dev Biol 275, 375–388. https://doi.org/10.1016/j.ydbio.2004.08.015.

Porpiglia, E., Samusik, N., Ho, A.T.V., Cosgrove, B.D., Mai, T., Davis, K.L., Jager, A., Nolan, G.P., Bendall, S.C., Fantl, W.J., et al. (2017). High-resolution myogenic lineage mapping by single-cell mass cytometry. Nat Cell Biol 19, 558–567. https://doi.org/10.1038/ncb3507.

Robinson, M.D., McCarthy, D.J., and Smyth, G.K. (2010). edgeR: a Bioconductor package for differential expression analysis of digital gene expression data. Bioinformatics 26, 139–140. https://doi.org/10.1093/bioinformatics/btp616.

Rocheteau, P., Gayraud-Morel, B., Siegl-Cachedenier, I., Blasco, M.A., and Tajbakhsh, S. (2012). A subpopulation of adult skeletal muscle stem cells retains all template DNA strands after cell division. Cell 148, 112–125. https://doi.org/10.1016/j.cell.2011.11.049.

Rodgers, J.T., King, K.Y., Brett, J.O., Cromie, M.J., Charville, G.W., Maguire, K.K., Brunson, C., Mastey, N., Liu, L., Tsai, C.R., et al. (2014). mTORC1 controls the adaptive transition of quiescent stem cells from G0 to G(Alert). Nature 510, 393–396. https://doi.org/10.1038/nature13255.

Sambasivan, R., Yao, R., Kissenpfennig, A., Van Wittenberghe, L., Paldi, A., Gayraud-Morel, B., Guenou, H., Malissen, B., Tajbakhsh, S., and Galy, A. (2011). Pax7-expressing satellite cells are indispensable for adult skeletal muscle regeneration. Development 138, 3647–3656. https://doi.org/10.1242/dev.067587.

Scaramozza, A., Park, D., Kollu, S., Beerman, I., Sun, X., Rossi, D.J., Lin, C.P., Scadden, D.T., Crist, C., and Brack, A.S. (2019). Lineage Tracing Reveals a Subset of Reserve Muscle Stem Cells Capable of Clonal Expansion under Stress. Cell Stem Cell 24, 944–957.e5. https://doi.org/10.1016/j.stem.2019.03.020.

Schmidt, M., Schüler, S.C., Hüttner, S.S., von Eyss, B., and von Maltzahn, J. (2019). Adult stem cells at work: regenerating skeletal muscle. Cell Mol Life Sci 76, 2559–2570. https://doi.org/10.1007/s00018-019-03093-6.

Schüler, S.C., Liu, Y., Dumontier, S., Grandbois, M., Le Moal, E., Cornelison, D., and Bentzinger, C.F. (2022). Extracellular matrix: brick and mortar in the skeletal muscle stemcell niche. Front Cell Dev Biol 10. https://doi.org/10.3389/fcell.2022.1056523

Seale, P., Sabourin, L.A., Girgis-Gabardo, A., Mansouri, A., Gruss, P., and Rudnicki, M.A. (2000). Pax7 is required for the specification of myogenic satellite cells. Cell 102, 777–786. https://doi.org/10.1016/s0092-8674(00)00066-0.

Shyu, A.-B., Wilkinson, M.F., and van Hoof, A. (2008). Messenger RNA regulation: to translate or to degrade. EMBO J 27, 471–481. https://doi.org/10.1038/sj.emboj.7601977

Sousa-Victor, P., García-Prat, L., and Muñoz-Cánoves, P. (2022). Control of satellite cell function in muscle regeneration and its disruption in ageing. Nat Rev Mol Cell Biol 23, 204–226. https://doi.org/10.1038/s41580-021-00421-2

Strack, R.L., Hein, B., Bhattacharyya, D., Hell, S.W., Keenan, R.J., and Glick, B.S. (2009). A rapidly maturing far-red derivative of DsRed-Express2 for whole-cell labeling. Biochemistry 48, 8279–8281. https://doi.org/10.1021/bi900870u.

Szymczak, A.L., Workman, C.J., Wang, Y., Vignali, K.M., Dilioglou, S., Vanin, E.F., and Vignali, D.A.A. (2004). Correction of multi-gene deficiency in vivo using a single “self-cleaving” 2A peptide-based retroviral vector. Nat Biotechnol 22, 589–594. https://doi.org/10.1038/nbt957.

Tajbakhsh, S., Rocancourt, D., and Buckingham, M. (1996). Muscle progenitor cells failing to respond to positional cues adopt non-myogenic fates in myf-5 null mice. Nature 384, 266–270. https://doi.org/10.1038/384266a0.

van Velthoven, C.T.J., and Rando, T.A. (2019). Stem Cell Quiescence: Dynamism, Restraint, and Cellular Idling. Cell Stem Cell 24, 213–225. https://doi.org/10.1016/j.stem.2019.01.001.

von Maltzahn, J., Jones, A.E., Parks, R.J., and Rudnicki, M.A. (2013). Pax7 is critical for the normal function os satellite cells in adult skeletal muscle. Proc Natl Acad Sci USA 110, 16474–16479. https://doi.org/10.1073/pnas.1307680110.

Zammit, P.S. (2017). Function of the myogenic regulatory factors Myf5, MyoD, Myogenin and MRF4 in skeletal muscle, satellite cells and regenerative myogenesis. Semin Cell Dev Biol 72, 19–32. https://doi.org/10.1016/j.semcdb.2017.11.011.

Zismanov, V., Chichkov, V., Colangelo, V., Jamet, S., Wang, S., Syme, A., Koromilas, A.E., and Crist, C. (2016). Phosphorylation of eIF2α Is a Translational Control Mechanism Regulating Muscle Stem Cell Quiescence and Self-Renewal. Cell Stem Cell 18, 79–90. https://doi.org/10.1016/j.stem.2015.09.020.

